# Adaptive Bayesian localization of motor representation areas

**DOI:** 10.64898/2026.06.01.729093

**Authors:** Mikael Laine, Tuomas P. Mutanen, Shokoofeh Parvin, Ole Numssen, Konstantin Weise, Matti Stenroos, Ida Granö, Ana M. Soto, Renan H. Matsuda, Victor H. Souza, Thomas R. Knösche, Risto J. Ilmoniemi

## Abstract

Transcranial magnetic stimulation (TMS) enables non-invasive localization of cortical motor representations, with important clinical applications in presurgical planning. Existing methods either disregard spatial information about the TMS-induced electric field (E-field) or use acquisition schemes that do not leverage previously elicited motor responses to guide subsequent stimulation. We present an adaptive Bayesian localization method that combines real-time E-field optimization with per-trial probabilistic inference. The cortical origin of the motor responses is represented as a spatial probability distribution that is updated after each stimulus. Subsequent stimuli are then optimized to maximize the expected localization improvement given previous responses. We validated the method experimentally, using multi-locus TMS for adaptive localization and single-coil TMS as a non-adaptive reference with randomized coil placements. Across eight subjects, the adaptive protocol at least halved the number of stimuli required for stable localization compared to the randomized protocol, converging in 60 stimuli on average, with 95% highest-density regions often below 10 mm^2^ by 150 stimuli.

## Introduction

Understanding the brain’s functional organization is both a fundamental scientific question and a matter of considerable clinical relevance for the treatment of brain disorders. Transcranial magnetic stimulation (TMS) provides a causal probe of this organization, in which, by stimulating the cortex non-invasively, one can elicit measurable responses such as motor evoked potentials (MEPs). TMS activates neurons by inducing an electric field (E-field) in the brain via a brief pulse of current driven through one or more coils placed on the scalp. Accurate localization of the cortical sites responsible for the stimulation-induced responses has clinical value, for example, in treatment planning for tumor resection and radiotherapy [1, 2], and in intraoperative identification of eloquent cortex [3] to avoid damaging healthy tissue. Greater localization accuracy could improve surgical outcomes, e.g., by minimizing craniotomy size, while shortening the mapping procedure time would reduce operational costs, patient discomfort, and in some cases, seizure risk [4].

Existing TMS localization methods differ in how the induced E-field is represented, how the stimulation-response relationship is modeled, and how stimulation sites are selected. The simplest representation of the E-field’s effect is to assume the activated cortex to be situated directly beneath the coil center [5] or at the E-field’s maximum or centroid [6, 7, 8], disregarding the E-field’s broader effect in activating the cortex. Recently, several methods that utilize the E-field distribution and find spatial correspondence between the E-field and the MEP amplitudes or motor thresholds have been proposed to improve localization [9, 10, 11, 12, 13, 14]. These approaches are based on the physiological assumption that, when TMS is delivered across multiple locations and intensities, MEP amplitude increases with E-field strength specifically at the true cortical representation site. Some approaches explicitly model the MEP amplitude as a non-linear function of the E-field strength [9, 12]. By requiring that the measured responses fit a specific motor input–output model, this stronger prior penalizes regions where the expected behavior is not observed. By applying an MEP amplitude regression model, high localization precision has been achieved with a fast, open-loop protocol in which coil placements are selected randomly over the motor cortex [15, 16]. Schultheiss et al. [17] subsequently combined this regression approach with prospective optimization of coil placements, reporting a near halving of the number of stimuli compared to random coil placements. While such open-loop acquisition keeps computation and data analysis outside the TMS session, which can allow for more careful planning and is robust against noisy observations, it cannot adaptively concentrate the stimuli in the most informative regions as MEP observations accumulate, resulting in more trials needed to achieve stable localization than a closed-loop approach would require.

In this work, we propose an adaptive Bayesian localization method that reduces the number of stimuli needed for localization of motor representation areas through real-time E-field optimization and simultaneously improves precision through probabilistic measurement interpretation. The cortical origin of MEPs is represented as a spatial probability distribution that is updated after each stimulus, and the E-field of each subsequent stimulus is optimized to maximize the expected improvement in localization. Unlike the probabilistic approach described in [18], which operates on motor thresholds, a summary statistic of many trials, our model uses individual stimulation trials, allowing the extraction of more information from the measurement data. Critically, the proposed model can estimate how the E-field direction modulates the response probability [19, 20], a crucial factor previously ignored [21, 22]. The probabilistic formulation has several benefits: it gives an intuitive measure of localization confidence, enables a principled stopping criterion based on achieved precision, and can incorporate prior beliefs about the probable location of the cortical origin of stimulation-induced responses.

For the adaptive protocol to be practical, E-field optimization, stimulus delivery, and inference must be realized fast enough so that the gains from reducing trial numbers are not lost to per-trial computational overhead. We achieve this using a multi-locus TMS (mTMS) system [23, 24], which controls the induced E-field electronically without physical coil movement, together with an open-source ABL (Adaptive Bayesian Localization) toolkit [25] that performs optimization and Bayesian inference in less than one second per trial, resulting in a fully automated protocol. We compare the adaptive protocol against a randomized figure-of-eight coil positioning similar to [15], and evaluate Bayesian inference against an MEP amplitude regression model based on the established method in [16]. In addition to the multi- and single-coil TMS demonstrated in this work, the proposed localization method could also be used, e.g., in direct cortical stimulation, or in general applications where a field can evoke binary responses that originate from a region.

## Methods

### Bayesian localization model

The probabilistic localization model identifies the cortical region whose E-fields across stimuli are most consistent with the observed responses, such as MEPs. The cortical ROI is partitioned into *N*_*q*_ non-overlapping regions 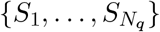 of approximately equal area, each centered on a vertex *q*. The E-field vector at vertex *q* on trial *t* is denoted by **E**_*q,t*_ *∈* ℝ^3^. Across *N*_*t*_ stimulation trials, the cortical E-fields 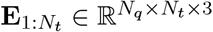 and the corresponding motor responses 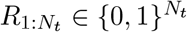 are obtained, with *R*_*t*_ = 1 assigned when a stimulation-evoked response is observed, e.g. when the response amplitude exceeds a chosen threshold *T*_*R*_.

The probability that the cortical origin of the responses lies in region *S*_*q*_ is denoted by *P* (*S*_*q*_), with a uniform prior *P* (*S*_*q*_) = 1*/N*_*q*_ inside the ROI and zero outside (Figure 1A). The trial-wise response probability is modeled as a modified sigmoid of the E-field strength |**E**_*q,t*_| and the E-field angle 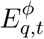. The latter is a scalar describing the orientation of **E**_*q,t*_ within a planar surface that captures the dominant E-field variation, measured as the in-plane angle relative to any chosen reference direction (see the *Implementation* section for its calculation). The response function is parametrized by *θ*_*q*_ = (*γ, δ, α*_1_, *α*_2_, *β*_1_, *β*_2_) and defined as

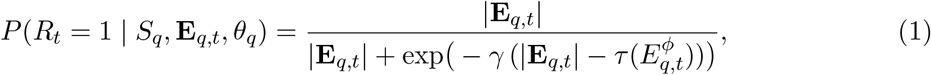

with the orientation-dependent excitability

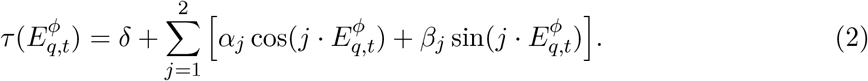

**Figure 1:**
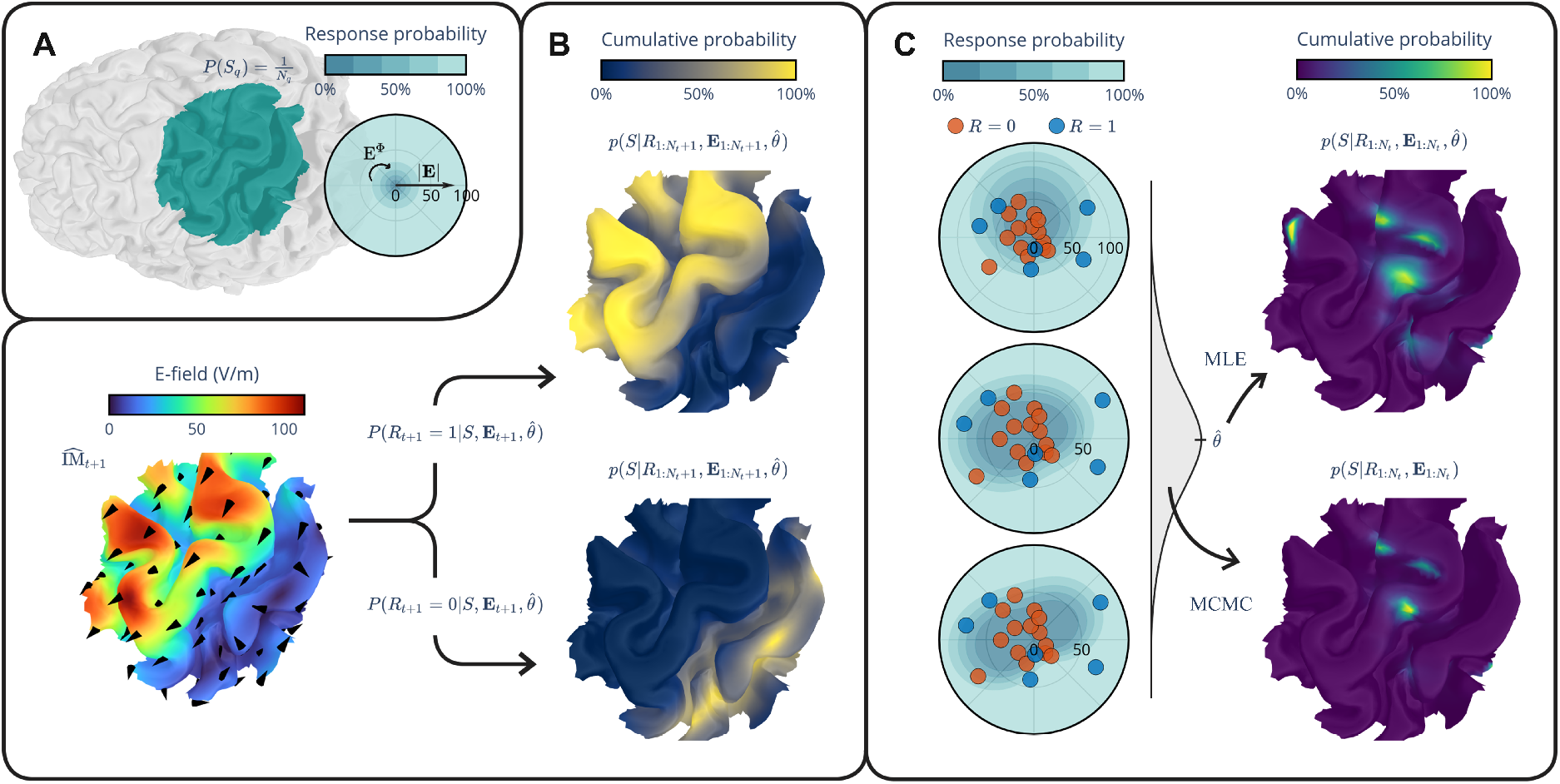
Principles of the adaptive Bayesian localization method. Notation is defined in the *Bayesian localization model* section, with an adaptation from vertex-based notation to distributions by omitting subscript *q* and substituting *P* with *p*. **A**. A cortical region of interest (ROI) is defined, and prior probabilities *P* (*S*_*q*_) and initial response function parameters *θ*_*q*_ are set for each region in the ROI. **B**. The E-field is optimized to find the maximum expected improvement 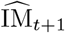 over the binary response *R*_*t*+1_. **C**. Using the acquired observations, the response function parameters are updated either by maximum likelihood estimation (MLE) to obtain point estimates, or by the more computationally demanding Markov chain Monte Carlo (MCMC) to obtain parameter distributions. The cortical origin of the observed responses is then identified using Bayes’ theorem.

The steepness of the sigmoid is controlled by *γ*, and the response curve is shifted along the E-field strength axis by *δ*. An orientation-dependent shift of the response curve is introduced by the Fourier coefficients *α*_*j*_ and *β*_*j*_ (*j* = 1, 2; see Figure 1C for response function visualizations). The probability of no response follows as *P* (*R*_*t*_ = 0 |*S*_*q*_, **E**_*q,t*_, *θ*_*q*_) = 1 − *P* (*R*_*t*_ = 1 |*S*_*q*_, **E**_*q,t*_, *θ*_*q*_). Because responses cannot be evoked without stimulation, the response probability must vanish as |**E**_*q,t*_| approaches zero, a constraint not satisfied by the standard logistic form. The chosen form of 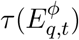 encompasses ellipse-like shapes and permits directional asymmetry: equal E-field strengths applied in opposite directions can yield different MEP probabilities, consistent with the direction-dependent asymmetry of MEP amplitudes reported in [20]. The directional dependence of the steepness parameter *γ* is not modeled, for simplicity.

Localization proceeds by updating the prior probability distribution according to the observations. Assuming conditional independence of trials, Bayes’ theorem gives

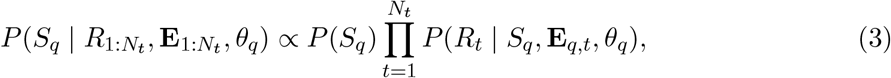

with the posterior distribution obtained by normalization: 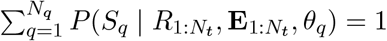.

Two inference approaches were used, differing in the treatment of the response function parameters *θ*_*q*_. During the automated measurement, a regularized maximum likelihood estimate (MLE) 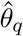 was computed in real time, and the posterior was conditioned on this point estimate:

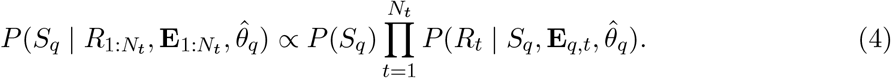

As a post-processing step, uncertainty in *θ*_*q*_ was accounted for by averaging the conditional posterior 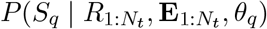 over the posterior of *θ*_*q*_, which was sampled via Markov chain Monte Carlo (MCMC):

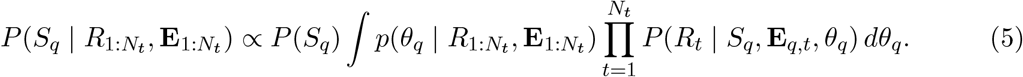

Results are reported using the MCMC posterior unless the MLE is explicitly specified.

The localization result is visualized as a spatial cumulative posterior: the regions are sorted in ascending order of posterior probability and, for each *S*_*q*_, the probabilities of all regions ranked at or below it are summed. Consequently, the least probable region receives a value near 0% and the most probable region receives 100%. Credible regions, such as the 95% highest-density region (HDR), can then be read directly from this representation as the set of regions for which the cumulative probability exceeds 5%. The 95% HDR area is approximated by assigning to each vertex one third of the area of every incident face and summing these vertex areas over all vertices inside the HDR.

### E-field optimization

Before each new trial *t* + 1, the E-field to be delivered can be optimized based on the current posteriors *P* (*S*_*q*_ | *R*_1:*t*_, **E**_1:*t*_, 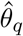) and the corresponding parameter estimates 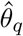, for all *q ∈* {1, …, *N*_*q*_}(Figure 1B). The objective of the optimization is to maximize the expected change in the posterior over the next trial, defined as

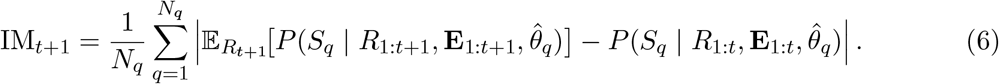

### MEP amplitude regression

The Bayesian approach was compared against an MEP amplitude regression approach based on element-wise regression of MEP amplitudes as a function of E-field strength, as in [16]. However, the regression fit was performed with a custom piecewise-linear function consisting of a horizontal baseline and a linear segment with a non-negative slope constraint. This customization was done because the generated motor input–output curves were limited to the initial portion of the expected sigmoidal curve, which was not well fitted by either a sigmoidal or a simple linear function (see Figures S2 and S3 in the Supplementary Material). The regression was performed with a custom Python script, following the same MLE routine used for the response function parameter fitting. Localization quality was assessed using the *R*^2^ goodness-of-fit statistic [16], which quantifies how well the regression model predicts the MEP amplitudes, with higher values indicating a better fit. Negative *R*^2^ values were clamped to zero, as they reflected poorly fitted models.

### Experimental validation

The proposed adaptive Bayesian localization method was validated in an experiment consisting of two measurements: one in which the mTMS system was used to localize a motor representation area with per-trial optimized E-fields (adaptive protocol), and a second in which stimulation was delivered with randomized figure-of-eight coil positioning over the motor cortex (randomized protocol).

### Implementation

Prior to the TMS sessions, an E-field model (see the *E-field modeling* section) was used to determine a pose (position and orientation) of the mTMS coil array, tangential to the scalp, at which its two figure-of-eight coils induced an E-field peak near the center of the hand knob on the cortical ROI. During adaptive localization, the coil-wise induced E-fields from this pose were used to optimize the E-field of every new trial (see the *E-field optimization* section):

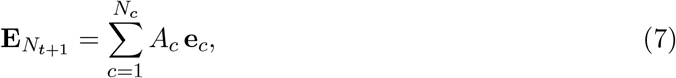

where *N*_*c*_ is the number of coils, *A*_*c*_ is the optimization parameter that represents the amplitude of the rate of change of current in coil *c*, and 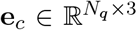 is the E-field of the *c*th coil given a unit rate of change of current. Three constraints were applied to the E-field optimization: First, the values *A*_*c*_ were bounded by the maximum output of the mTMS device. Second, the maximum E-field strength in the cortex was limited to 200 V/m to avoid subject discomfort due to excessive E-field on the scalp. Third, the mTMS pulse was constrained not to exceed the mechanical strain limit specified by the mTMS control software (see *mTMS pulse delivery and strain limit* section in the Supplementary Material). E-field optimization was performed using a differential evolution algorithm from the evosax library [26]. Although the adaptive protocol in this work used the mTMS system, the E-fields could also be optimized for single-coil TMS, e.g., by using a precomputed library of E-fields corresponding to different coil poses.

The E-field direction at each ROI vertex was summarized as a scalar angle *E*_*ϕ*_ to simplify its interpretation in the response function. In a spherically symmetric model, volume conductor effects would lead to a zero radial field component regardless of coil position and orientation. In a realistically shaped conductor, for a given field point, this phenomenon manifests as the E-field vector varying primarily in only two dimensions. Therefore, the set of E-field vectors at each vertex was individually projected onto a plane that preserved the most variance, identified by principal component analysis (PCA). For mTMS, PCA was applied across the five distinct E-field patterns of the coil array. For the Magstim coil, PCA was applied across the full set of E-fields induced during the localization procedure. Fixed reference directions where *E*_*ϕ*_ = 0^*°*^ were arbitrarily chosen by projecting the positive *y*-axis of the coordinate system onto each vertex’s plane. *E*_*ϕ*_ was then computed as the angle between the projected E-field vector and the reference direction.

To fully automate the adaptive Bayesian localization measurement on the mTMS hardware, an open-source Python toolkit, ABL [25], was developed to handle the E-field optimization and inference, and a custom MATLAB script was used to deliver the optimized E-field and record the MEPs on the mTMS system. The MEP detection threshold *T*_*R*_ was set to 30 *µ*V, approximately three times the mean peak-to-peak baseline electromyography (EMG) noise of the filtered signal (10.2 *±* 3.9 *µ*V, mean *±* SD, measured across subjects, muscles, and trials). The measurement data from the randomized protocol were also analyzed post-experimentally using the ABL toolkit. The Bayesian probability distributions were evaluated with vectorized operations using the PyTorch library [27]. The MLE coefficients 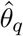 were obtained with a gradient-based Adam optimizer, and MCMC sampling was performed with Pyro [28].

To assess the computational demands of the adaptive protocol, per-trial execution times were measured by reproducing all experiments in simulation on a benchmark system equipped with an NVIDIA RTX A2000 GPU (4 GB VRAM) and an Intel Core i7-11850H CPU, allowing a direct comparison between GPU- and CPU-based execution. Four quantities were timed: a one-time precompilation step required before the first trial, E-field optimization, MLE inference, and MCMC inference.

### E-field modeling

Fat-suppressed T1- and T2-weighted magnetic resonance images (MRI) were acquired using a 3T MAGNETOM Skyra scanner (Siemens Healthcare, Erlangen, Germany) at the Advanced Magnetic Imaging (AMI) Centre, Aalto NeuroImaging, Aalto University School of Science. The induced E-fields were computed using reciprocal Galerkin boundary element method as described in [29]. The Magstim 70 mm Double Coil (P/N 9925; Whitland, United Kingdom) was modeled using a set of magnetic dipoles as in [29], and the mTMS coils by discretizing the coil windings into sets of current dipoles as in [23]. The head model was prepared using the SimNIBS mri2mesh pipeline [30]. The E-field was evaluated at the mid-cortical layer segmented with FreeSurfer [31], within a cortical ROI of approximately 5 cm in diameter centered on the left hand knob [32]. The ROI had approximately 5000 vertices and a vertex-to-vertex distance of 1.2 mm.

### TMS–EMG

Eight right-handed subjects (5 female, 3 male; aged 27–36 years) participated in the experiment. The study was approved by the Ethics Committee of the Hospital District of Helsinki and Uusimaa (HUS/1198/2016) and conducted in accordance with the Declaration of Helsinki. The subjects gave their informed consent in writing. At the start of the measurement, subjects were seated on a reclining chair and instructed to avoid movement and to keep their right hand relaxed during stimulation. During stimulation, subjects wore hearing protection.

For the EMG recording (NeurOne; Bittium Plc, Finland), surface electrodes (wet gel; Ag/AgCl; Spes Medica, Italy) were placed on the right abductor pollicis brevis (APB), first dorsal interosseous (FDI), and abductor digiti minimi (ADM) muscles in a belly-tendon montage. The ground electrode was attached to the ulnar styloid process. EMG of the FDI was monitored during stimulation, and if muscle preactivation was observed, subjects were instructed to relax their hand. During adaptive localization, EMG segments recorded from the FDI muscle, spanning 200 ms before to 100 ms after stimulation, were extracted and subsequently processed using a 20–500 Hz band-pass filter and a 50 Hz spectral interpolation filter [33] to remove line noise. For each EMG segment, the MEP peak-to-peak amplitude was calculated from a time window of 15–45 ms after the stimulus.

TMS was delivered with two systems in a randomized order of application: a Magstim 70 mm Double Coil connected to a Magstim 200^2^ (Magstim Co Ltd, UK), and a five-coil mTMS array connected to our custom power electronics [23, 24]. For the Magstim system, a sinusoidal monophasic waveform was used, whereas for the mTMS five-coil array, a trapezoidal monophasic waveform was used.

With the Magstim system, 150 stimuli were applied with randomized coil positioning over the motor cortex (randomized protocol). The coil position was varied within a region approximately 5 cm in diameter centered on the hand knob, and the coil was rotated around the scalp normal within an approximately 180^*°*^ range centered in the anteromedial direction. The stimulation intensity was set after a brief motor mapping to ensure that the stimulation could produce MEP amplitudes above 200 *µ*V.

The mTMS system delivered 150 stimuli using the adaptive localization method for the FDI muscle (adaptive protocol), with a 5-minute coil-cooling break introduced midway through the measurement. The coil array was maintained at the predetermined pose via robotic control [34, 35] with an Elfin 5 (Han’s Robot Co Ltd, China), with a maximum of 2 mm or 3^*°*^ of error in location and direction, respectively. InVesalius neuronavigation software [36] with individual MRIs was used together with eight Flex 13 infrared cameras (OptiTrack; NaturalPoint Inc, USA) for tracking the coil and the subject’s head. MRI co-registration was performed using a refinement consisting of approximately 50 points. The inter-stimulus interval was randomized between 2 and 3 seconds via software control with the mTMS system. With the Magstim system, a similar interval was maintained using pedal triggering.

### Post-processing

The induced E-fields from the randomized protocol were modeled following the TMS session. When the recorded coil poses intersected the scalp (e.g., due to head co-registration error), they were translated along the scalp normal to avoid the intersection while maintaining the coil orientation. The resulting translation was 0.4 *±*0.6 mm (mean *±*SD) across subjects and coil poses. The rate of change of current was derived by scaling the subject-specific stimulator output percentage to the device’s maximum output of 114.7 A/*µ*s [37].

The recorded EMG from both measurement protocols was post-processed to obtain MEP amplitudes. Processing was equivalent to that in the online recording, except that the band-pass filter was applied to the whole recording and its high-pass cutoff was changed to 10 Hz to reduce temporal smearing. The EMG trials were screened for preactivation by flagging trials where the MEP exceeded 20 *µ*V and the EMG signal’s peak-to-peak amplitude in the window from 120 to 20 ms before stimulus onset also exceeded 20 *µ*V. After visual inspection, flagged trials with clear preactivation were removed. The threshold of 20 *µ*V was chosen as a sensitive estimate of motor activity that remains distinguishable from baseline noise of approximately 10 *µ*V. For the APB, FDI, and ADM muscles, 1.7%, 0.7%, and 3.1% of MEPs were rejected due to preactivation, respectively.

## Results

### Adaptive Bayesian localization

Figure 2 shows the localization of APB, FDI, and ADM obtained with the Bayesian model under the adaptive protocol (see Figures S1, S2, and S3 in the Supplementary Material for localization results from the other method–protocol combinations). The cumulative probability peaks lie on the crown, rim, or sulcal wall of the precentral gyrus. Within each subject, the localization maps across muscles were similar, with localization peaks coinciding within 3 mm of each other. The estimated response functions indicated a pronounced effect of E-field direction on MEP probability, most evident in S3 and S4, which showed a figure-of-eight-shaped orientation dependence in all muscles (Figure 2). In several subjects, the sampled E-field directions clustered within narrow angular windows perpendicular to the course of the central sulcus, likely reflecting the intensity constraints imposed by the E-field optimization procedure.

**Figure 2:**
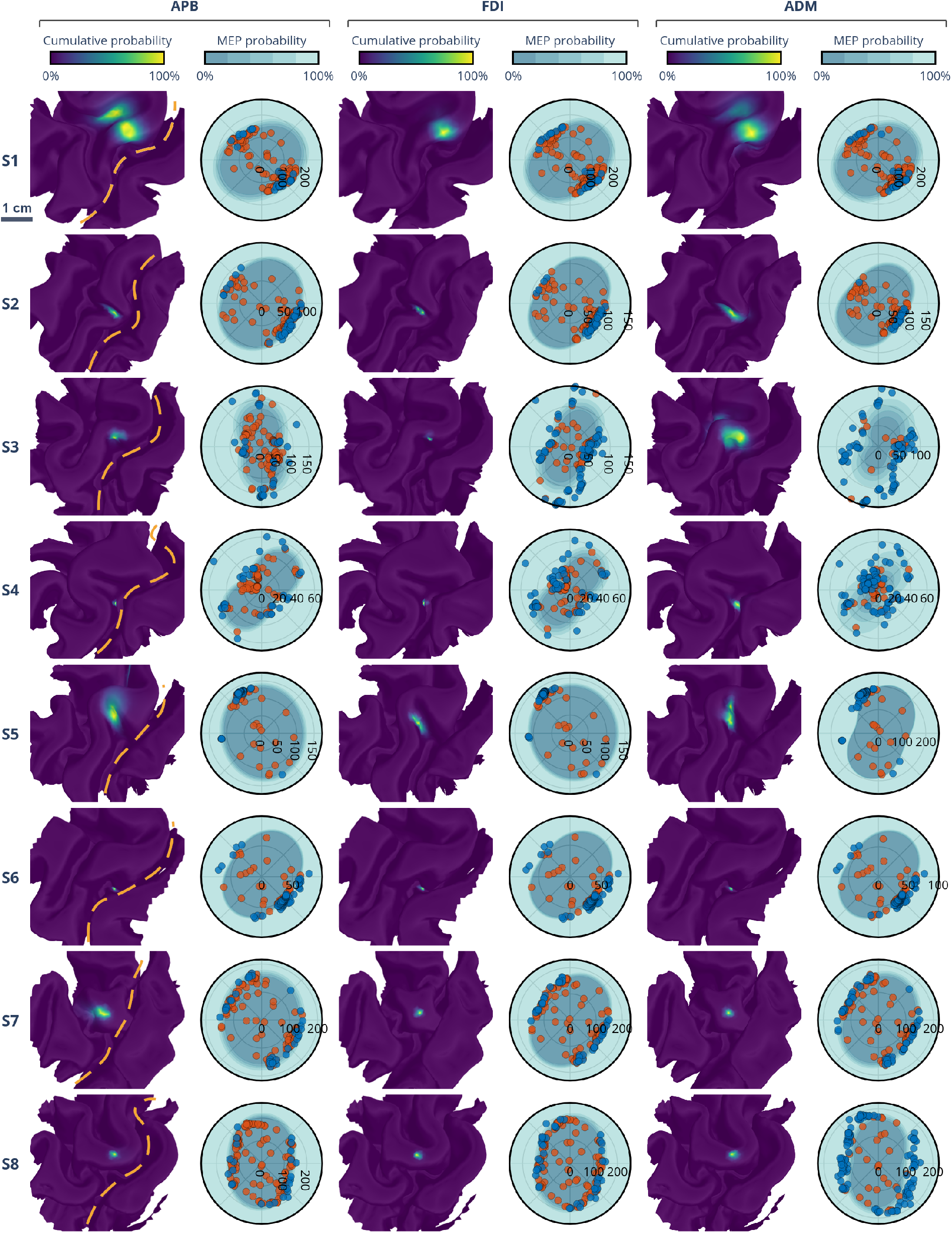
APB, FDI, and ADM muscle localizations with the Bayesian model under the adaptive protocol. Subjects S1–S8 are listed in rows. For each muscle, the cortical mesh on the left shows the localization map as a cumulative probability distribution, and the polar plot on its right shows the estimated response function at the most probable region. In the polar plot, the radial axis represents the E-field strength |**E**| and the angular axis the E-field angle *E*^*ϕ*^, which aligns with the view of the mesh. Each MEP is marked in the polar plot as a dot, colored blue if the MEP was detected (amplitude *>* 30 *µ*V) and red otherwise. The response function coefficients *θ*_*q*_ are averaged over their posterior distributions. The dashed line indicates the central sulcus.

The localization maps were generally precise, with the 95% HDR falling below 10 mm^2^ in many cases (Table 1). The 95% HDR was not strongly dependent on the number of detected MEPs; for example, in S8, APB and ADM reached similar 95% HDR values (10 mm^2^ and 13 mm^2^), while the numbers of detected MEPs were 25/150 and 120/150, respectively. Across the three muscles, FDI had the lowest 95% HDR in all but one subject, likely a result of optimizing the adaptive protocol specifically for FDI. The exception was S6, in which all three muscles had similarly low 95% HDR values, ranging from 2 mm^2^ to 4 mm^2^.

**Table 1:**
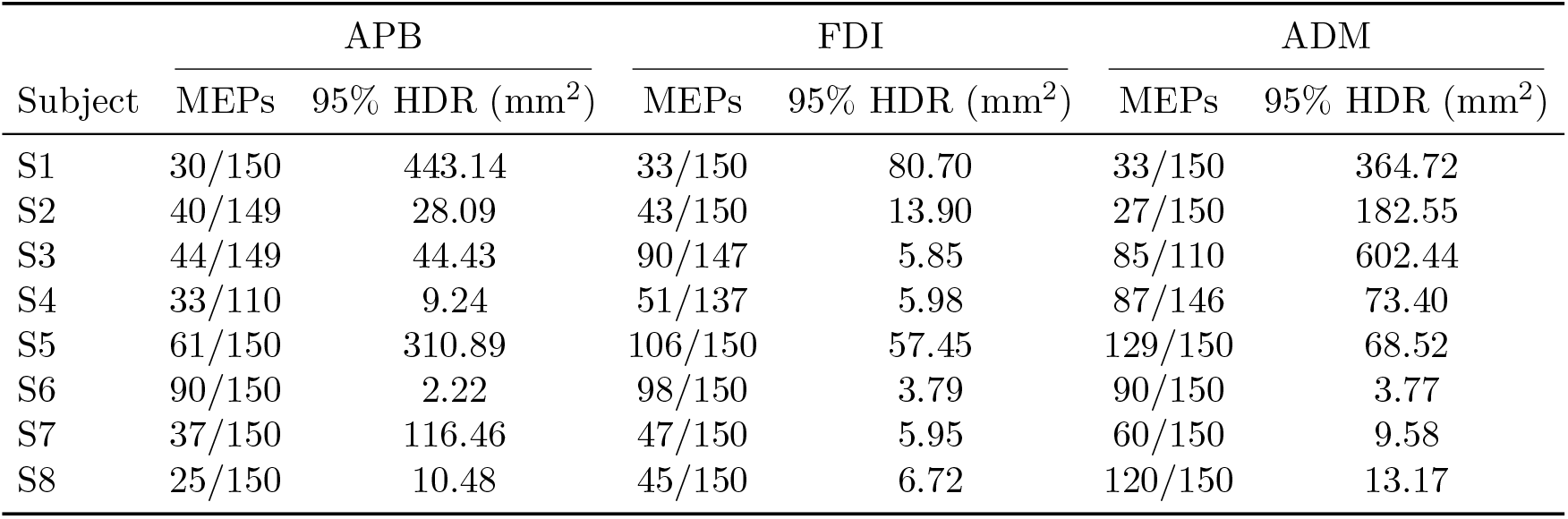
Statistics for localization maps of the Bayesian model under the adaptive protocol for subjects S1–S8. MEPs = the fraction of detected MEPs out of all trials; 95% HDR = 95% highest-density region.

| Subject | APB |  | FDI |  | ADM |  |
| --- | --- | --- | --- | --- | --- | --- |
|  | MEPs | 95% HDR (mm<sup>2</sup>) | MEPs | 95% HDR (mm<sup>2</sup>) | MEPs | 95% HDR (mm<sup>2</sup>) |
| S1 | 30/150 | 443.14 | 33/150 | 80.70 | 33/150 | 364.72 |
| S2 | 40/149 | 28.09 | 43/150 | 13.90 | 27/150 | 182.55 |
| S3 | 44/149 | 44.43 | 90/147 | 5.85 | 85/110 | 602.44 |
| S4 | 33/110 | 9.24 | 51/137 | 5.98 | 87/146 | 73.40 |
| S5 | 61/150 | 310.89 | 106/150 | 57.45 | 129/150 | 68.52 |
| S6 | 90/150 | 2.22 | 98/150 | 3.79 | 90/150 | 3.77 |
| S7 | 37/150 | 116.46 | 47/150 | 5.95 | 60/150 | 9.58 |
| S8 | 25/150 | 10.48 | 45/150 | 6.72 | 120/150 | 13.17 |

### Convergence and early stopping

The adaptive protocol converged substantially faster than the randomized protocol (Figure 3). Averaged across subjects, the localization peak stabilized within 2 mm of its final position after approximately 60 stimuli under the adaptive protocol, whereas under the randomized protocol the peak continued to shift by several millimeters beyond 120 stimuli, and the convergence curve showed no clear stabilization at 150 stimuli. The 95% HDR followed a similar pattern: under the adaptive protocol it contracted to 2 cm^2^ after 40 stimuli, whereas under the randomized protocol it remained above this value throughout the 150 stimuli. Together, these results indicate that adaptive E-field optimization at least halves the data requirement for stable localization.

**Figure 3:**
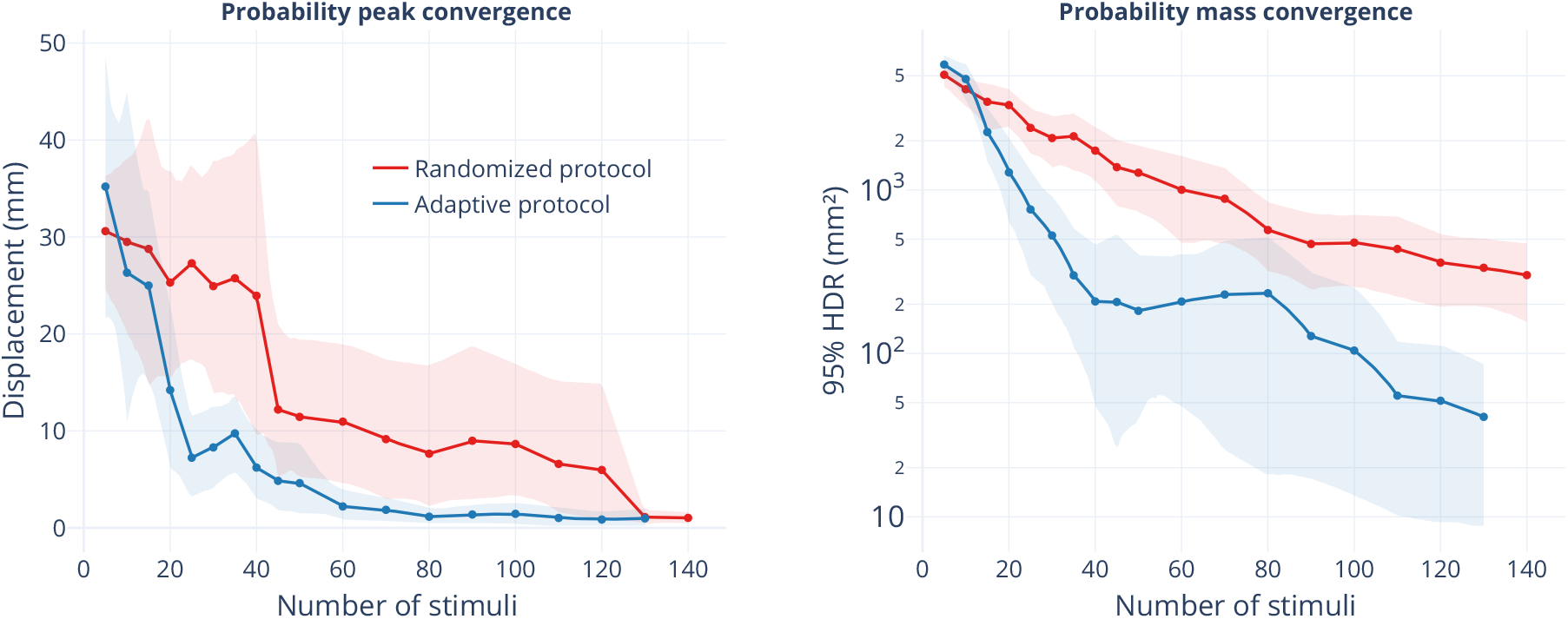
FDI localization convergence, averaged across subjects. The number of stimuli is shown up to the highest common number across subjects. Localization peaks are computed every 5 trials up to trial 50 and every 10 trials thereafter. Shaded regions indicate 95% confidence intervals. Left: geodesic displacement of the probability peak from its final position. Right: contraction of the probability mass, measured as the 95% HDR on a logarithmic scale.

Between 50 and 80 stimuli, the contraction of the 95% HDR briefly paused and slightly reversed. This transient plateau may reflect a shift in the adaptive algorithm from exploration toward exploitation: once the probability distribution concentrated on a single region, subsequent stimuli targeted the vicinity of that region and may occasionally have produced observations that were unexpected under the assumed posterior.

To evaluate the feasibility of early stopping during localization, we computed the localization convergence for the MLE posteriors, which are readily available during data acquisition, and defined a convergence criterion: localization was considered converged once the probability peak remained within a 5 mm region for 35 consecutive trials.

The selected criterion identified convergence without being triggered by momentary plateaus, with the exception of S1 under the randomized protocol, where convergence was falsely detected at a plateau at which the localization peak remained stable for around 50 stimuli before shifting by nearly 20 mm (Figure 4). The number of stimuli required to reach convergence ranged from 54 to 120 under the adaptive protocol and from 70 to 102 under the randomized protocol. However, S3, S5, and S8 never met the early stopping criterion under the randomized protocol. After post-processing the converged localizations with MCMC, the localization peaks were within 2 mm of the best estimates obtained with the full 150 trials.

**Figure 4:**
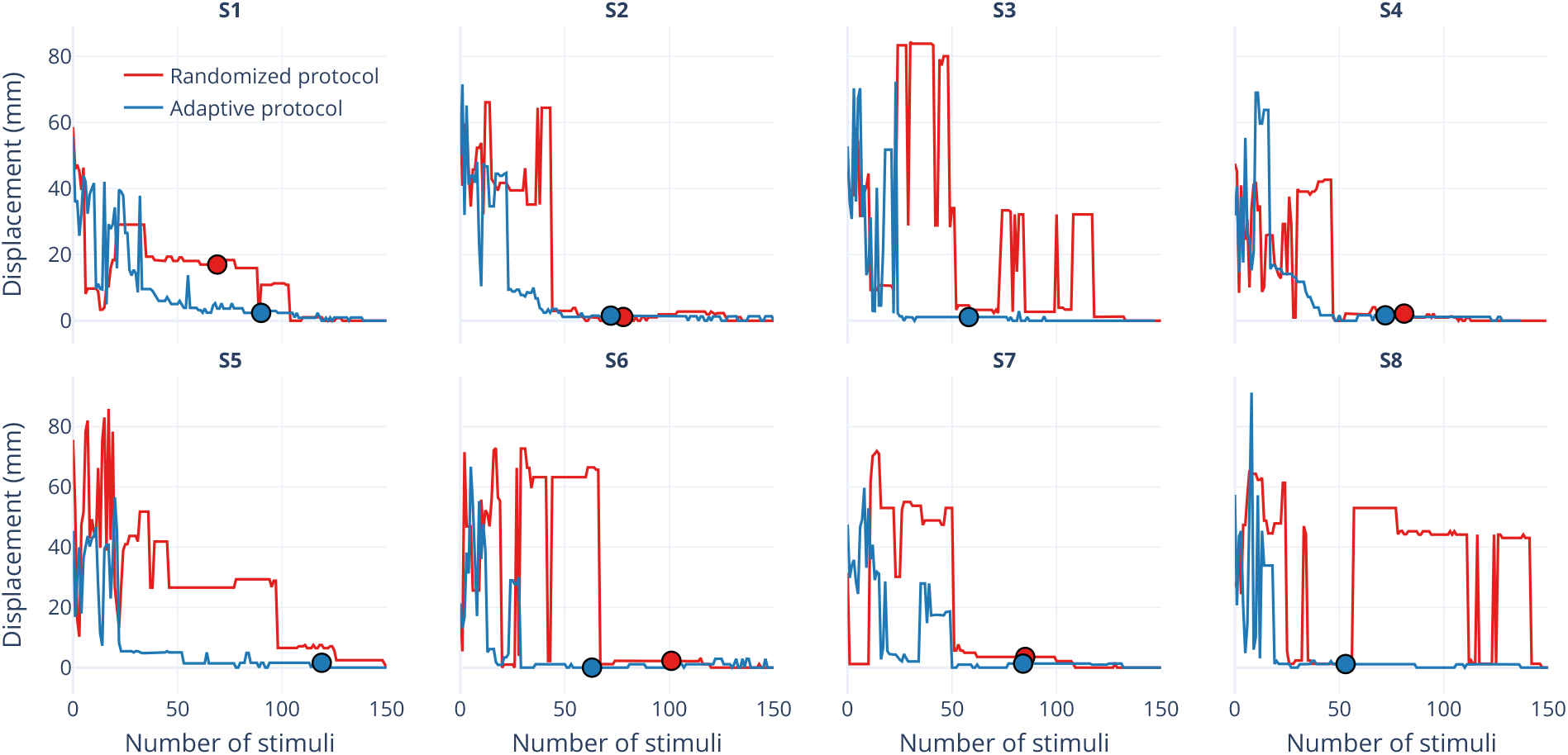
Geodesic displacement of the probability peak from its final position in each subject with MLE posteriors. The colored dots indicate when the early stopping criterion was met.

### Computation times

Use of the GPU enabled high computational speed with little delay between stimulation trials (Table 2). On the GPU, the per-trial time for E-field optimization and Bayesian inference with MLE was 0.6 s on average. MCMC inference on the GPU took 4 minutes, making it impractical for per-trial use but feasible for computing the final posterior within the measurement session. On the CPU, the per-trial computation time increased to 8.7 s, dominated by the 8.4 s E-field optimization, while MCMC inference took 48 minutes on average, with considerable variability.

**Table 2:** Computation times of E-field optimization and Bayesian inference on a benchmark system (see the *Methods: Implementation* section). Values represent the mean *±*SD. Precompilation times are averaged across subjects, E-field optimization and Inference (MLE) times over subjects and trials, and Inference (MCMC) times over subjects and select trials: every 5 trials up to trial 50 and every 10 trials thereafter.

| Device | GPU | CPU |
| --- | --- | --- |
| Precompilation | $3.30 \pm 0.18$ s | $9.45 \pm 1.98$ s |
| E-field optimization | $0.40 \pm 0.11$ s | $8.39 \pm 2.04$ s |
| Inference (MLE) | $0.18 \pm 0.10$ s | $0.27 \pm 0.11$ s |
| Inference (MCMC) | $3.65 \pm 1.73$ min | $47.80 \pm 36.49$ min |

### Localization comparison across methods and protocols

Figure 5 shows the FDI localization maps for all subjects, separated by inference method and protocol, and Table 3 provides additional statistics on the MEP detection rates, 95% HDRs, maximum MEP amplitudes, and maximum *R*^2^ values. Because the two methods estimate fundamentally different quantities, their spatial extents are not directly comparable across methods. We therefore compared spatial extent only between protocols within a given model, and compared the two methods through their map peaks.

**Figure 5:**
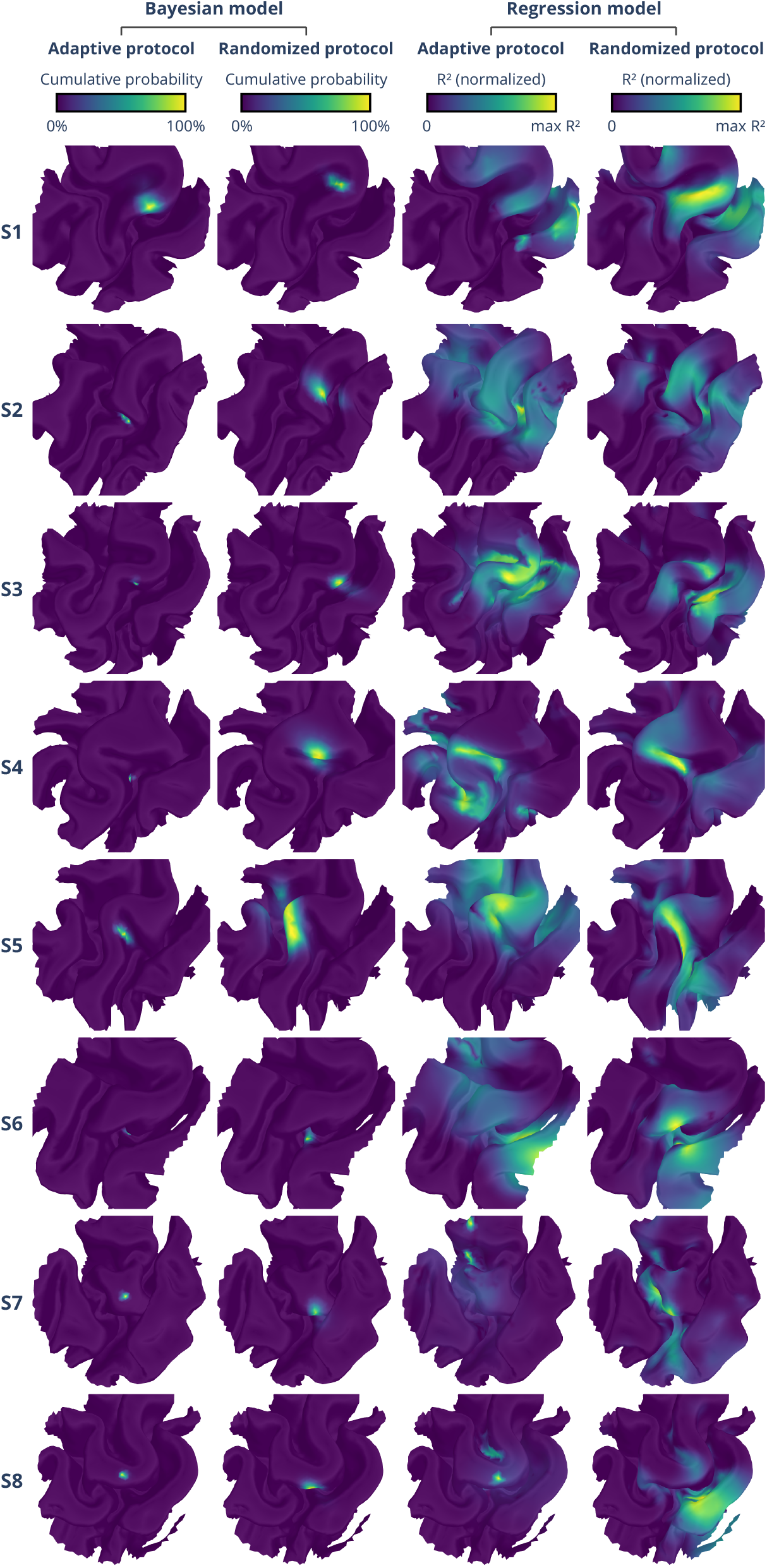
FDI localization map comparison across methods and protocols. Subjects S1–S8 are listed in rows. The first and second columns show the cumulative probability distributions from the Bayesian model for the adaptive and randomized protocols, respectively. The third and fourth columns show the normalized *R*^2^ maps from the MEP-amplitude regression approach for the same two protocols.

**Table 3:** Statistics for localization maps of the four method–protocol combinations for subjects S1–S8. MEPs = the fraction of detected MEPs out of all trials; 95% HDR = 95% highest-density region; MEP max = maximum MEP amplitude; max *R*^2^ = maximum *R*^2^ value across the map.

| Subject | Bayesian / adaptive |  | Bayesian / randomized |  | Regression / adaptive |  | Regression / randomized |  |
| --- | --- | --- | --- | --- | --- | --- | --- | --- |
| | MEPs | 95% HDR (mm<sup>2</sup>) | MEPs | 95% HDR (mm<sup>2</sup>) | MEP max (mV) | max $R^2$ | MEP max (mV) | max $R^2$ |
| S1 | 33/150 | 80.7 | 89/150 | 41.7 | 2.72 | 0.22 | 7.45 | 0.71 |
| S2 | 43/150 | 13.9 | 54/150 | 358.3 | 1.62 | 0.12 | 2.08 | 0.21 |
| S3 | 90/147 | 5.8 | 50/150 | 173.6 | 3.78 | 0.79 | 1.82 | 0.54 |
| S4 | 51/137 | 6.0 | 45/150 | 277.5 | 10.32 | 0.93 | 1.88 | 0.39 |
| S5 | 106/150 | 57.8 | 31/150 | 832.6 | 6.32 | 0.10 | 2.23 | 0.35 |
| S6 | 98/150 | 3.8 | 76/150 | 54.8 | 5.25 | 0.16 | 3.66 | 0.51 |
| S7 | 47/150 | 6.0 | 51/150 | 189.7 | 1.15 | 0.29 | 2.80 | 0.37 |
| S8 | 45/150 | 6.7 | 36/150 | 157.8 | 1.08 | 0.51 | 0.46 | 0.37 |

Within the Bayesian model, the 95% HDRs were smaller under the adaptive protocol (4– 80 mm^2^) than under the randomized protocol (40–830 mm^2^) in seven of eight subjects, indicating more informative data acquisition under real-time optimization; the same trend was evident in the convergence behavior (Figure 3). Within the regression model, the localization maps showed some correspondence between the protocols, although in some subjects the localization peaks fell on separate gyri. MEP detection ratios and maximum MEP amplitudes varied across subjects and protocols but showed no systematic difference between the adaptive and randomized protocols (Table 3).

To compare localization across methods and protocols, we examined the displacement between map peaks. The four method–protocol combinations produced similar peaks in some subjects, but no consistent pattern was apparent across the approaches (Figure 5). To quantify agreement, we defined the cumulative probability peak of the Bayesian model under the adaptive protocol as a per-subject reference and measured the geodesic displacement of all other method–protocol combinations relative to it (Figure 6A). This reference does not necessarily correspond to the ground truth, so deviations from it should not be interpreted as localization errors. Localization peaks were closest to the reference for the Bayesian model under the randomized protocol, with a median displacement of 8 mm across subjects and muscles. The regression model showed greater, and more variable, peak displacement under the adaptive protocol than under the randomized protocol, with median displacements of 28 mm and 15 mm, respectively. Because MEP detection ratios and maximum MEP amplitudes were similar between the protocols, this counterintuitive result may instead reflect the broader range of E-field directions induced by adaptive optimization: in the randomized protocol, the coil orientation was varied within an approximately 180^*°*^ range, correspondingly restricting the induced E-field directions, whereas the adaptive protocol could induce arbitrary directions. This wider directional range may have influenced the MEP amplitudes in a manner not accounted for by the regression model.

**Figure 6:**
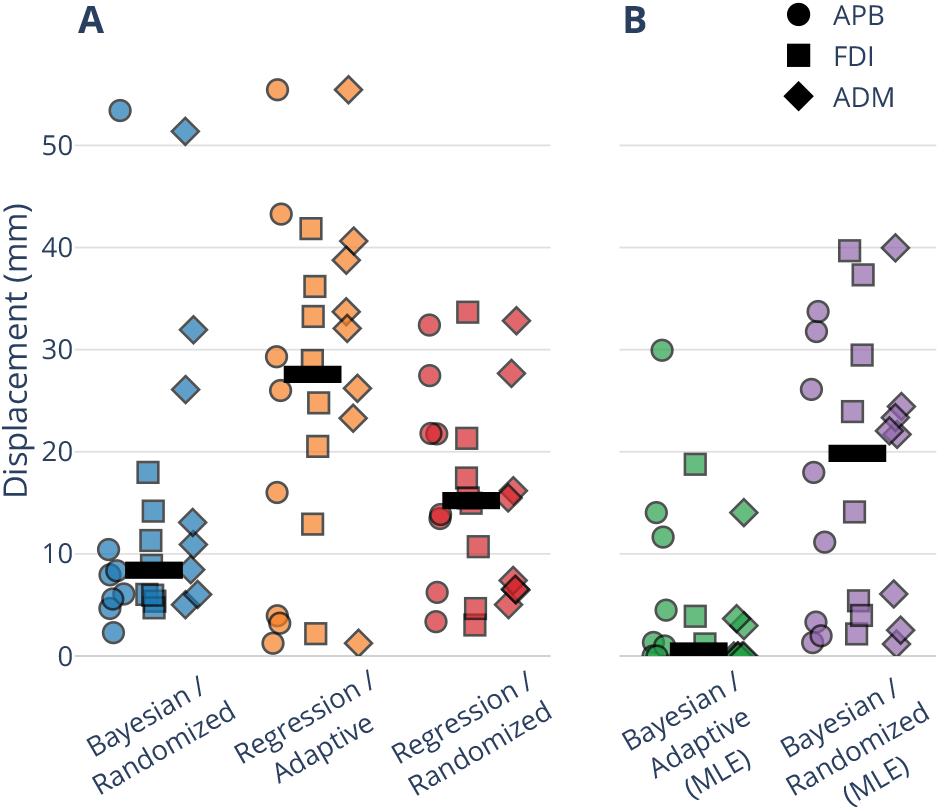
Comparison of geodesic localization peak displacements. The black horizontal bar in each group indicates the group median. **A**. Comparison across methods and protocols, with the Bayesian model (MCMC) under the adaptive protocol as a shared reference. **B**. Comparison of MLE posteriors against each protocol’s MCMC posterior.

Finally, we compared the localization peaks of the MLE posteriors with those of the corresponding MCMC posteriors (Figure 6B). Because the two variants share the same model but the MCMC posterior additionally integrates over parameter uncertainty, we treat the latter as the more accurate. The median displacement was 0 mm under the adaptive protocol: most of the time, the MLE posterior peak coincided with the MCMC peak. Under the randomized protocol, the median displacement was substantially larger, at 20 mm. The adaptive protocol may inadvertently constrain the response function parameters more tightly, resulting in a smaller benefit from the MCMC post-processing.

## Discussion

We present an adaptive Bayesian localization method for identifying cortical motor representations using TMS. The key advance over existing approaches is the combination of real-time E-field optimization and per-stimulus probabilistic inference, which together reduce the number of stimuli needed to reach a stable and precise localization. In addition, the localization employed a parametric model of the E-field orientation’s effect on the stimulation-induced responses. The Bayesian model produced focal posterior distributions concentrated on the crown, rim, or sulcal walls of the precentral gyrus (Figures 2 and 5), consistent with the known motor cortex organization [32, 38] and with the findings of existing localization approaches [9, 10, 11, 13, 18]. Furthermore, representations of different finger movements were consistently localized within 3 mm of each other, and the 95% HDR fell below 10 mm^2^ in many cases (Table 1). Averaged across subjects, the adaptive protocol at least halved the number of stimuli, and thus the measurement time, relative to the randomized protocol, with the localization peak stabilizing within 2 mm of its final value after 60 stimuli (Figure 3). The Bayesian model employed an approximate real-time inference, which could be used to stop localization after convergence, and a more robust post-processing inference that accounts for uncertainty of the modeled parameters.

### Efficiency and the role of E-field optimization

The reduction in the number of stimuli required to reach stable localization under the adaptive protocol versus the randomized protocol was comparable to the findings of [17], where prospectively optimized coil positioning nearly halved the required number of trials compared to randomized figure-of-eight coil positioning. Our adaptive method exceeds this efficiency while providing the practical advantages of not requiring any prior stimulation (e.g., motor threshold estimation) and of delivering stimuli without additional delay from coil repositioning when using the mTMS device [23, 24]. However, the Bayesian inference component is decoupled from the mTMS hardware and can be applied to data collected with conventional figure-of-eight coils, as our analysis of the Magstim measurements demonstrates.

Using an early stopping criterion based on localization peak stability could further reduce the measurement time, with stabilization reached within 50–120 stimuli under the adaptive protocol (Figure 4). With the randomized protocol, however, applying a convergence criterion was less reliable: convergence was falsely detected in one subject, while in three other subjects it was not reached within 150 stimuli. Future development may improve early stopping detection, e.g., by estimating the uncertainty of the response function parameters.

Real-time adaptive operation in the present setup benefited greatly from GPU acceleration: per-trial computation completed in 0.6 s on average on the GPU but increased to nearly 9 s on the CPU, with the E-field optimization accounting for most of the CPU cost (Table 2). Final MCMC posterior estimation took minutes on the GPU and nearly one hour on the CPU. The present method is therefore feasible to perform on standard hardware; however, without GPU acceleration, the post-processed localization may not be available within the localization session. The developed ABL toolkit [25] supports both CPU and GPU computations.

### Precision

The Bayesian posteriors were spatially concentrated on a single, focal region of the motor cortex (Figure 2). The final per-subject 95% HDR areas, reaching around 10 mm^2^ (Table 1), were similar in scale to the 95% HDR volume limit of around 200 mm^3^ reported in [18]: if these HDR regions were treated as a square and a cube, their side lengths would be 3.2 and 5.8 mm, respectively. The achievable precision may be referred to as super-resolution, since, by compounding information across trials, the precision is well below the effective extent of the induced E-field at cortical depth (full-width-at-half-maximum on the order of a few cm; see, e.g., [39]).

### Comparison with MEP amplitude regression

Agreement between the Bayesian and MEP amplitude regression localization peaks varied across subjects, and the pattern of identified representation regions was not consistent across all subjects (Figures 5 and 6). Our stimulation protocols elicited MEPs roughly half of the time (Table 3), producing motor input–output curves limited to the lower portion of the sigmoidal range, whereas the regression approach has been demonstrated and validated for a protocol using a stimulation intensity of around 150% of resting motor threshold and a biphasic waveform [16]. The use of monophasic pulses and lower intensities likely degraded regression performance relative to its intended operating regime. However, the ability to perform localization at a low intensity can be beneficial for patient safety and protocol tolerability.

Despite these across-method challenges, similar representation regions were identified in some subjects. A definitive evaluation of the developed method’s accuracy, however, requires an independent ground truth such as direct electrical stimulation [40].

### Limitations

Two observations point to limitations of the current Bayesian model. First, the posteriors from the adaptive and randomized protocols within the same subject did not coincide (Figures 5 and 6). This discrepancy could reflect a systematic error in the measurement data (e.g., coil placement or the model of the coil windings) or a genuine physiological effect of the measurement protocol, such as E-field orientation affecting which cortical region is responsible for an observed MEP [41, 42, 43]. Simulations with realistic neuronal morphologies suggest that an anteriorly directed E-field best activates neurons on the posterior rim of the precentral gyrus, whereas a posteriorly directed E-field best activates the anterior rim [41]. Indeed, the generally anteromedially directed E-fields of the randomized protocol produced localization peaks at the posterior rim in five out of eight subjects, compared to three under the adaptive protocol (Figure 5). Second, the response function fitting under the adaptive protocol used MLE point estimates, which disregard parameter uncertainty, leading to overconfidence in the probability posterior. Consequently, the localization peak can be less stable across stimuli, and the E-field optimization performance can degrade if an overconfident posterior leads to less effective exploration of the cortical ROI.

In addition to the response function parameter estimation, several modeling assumptions may contribute to overconfidence in the current framework. The observation likelihood assumes a single cortical source and a particular parametric form for the response function, and it treats both the induced E-fields and the measured MEPs as noise-free. In practice, neuronavigation error, head-model approximations, and hardware-related errors introduce E-field uncertainty that is not propagated into the posterior. Adding an E-field noise model and relaxing the single-source assumption are both natural extensions and would be expected to prevent overconfidence. A further limitation is the binary detection criterion: by thresholding at 30 *µ*V, the model discards information about the MEP amplitude. A response model based on a full MEP amplitude distribution rather than on detection probability could incorporate the physiological prior used in the MEP amplitude regression approach.

### Future directions

Several extensions of the presented method are possible. The E-field optimization is one-step optimal and maximizes the expected improvement of the next trial, whereas an *n*-step lookahead could plan informative stimulation sequences and may mitigate the brief plateau in 95% HDR contraction that we observed between 50 and 80 stimuli. A multi-muscle objective, summing the expected information gain across target muscles, would enable more effective joint localization of muscle groups. Relaxing the assumption of a single source region would enable a better understanding of the physiological processes related to response generation and could expand the applications of TMS as a neuroimaging tool.

In summary, combining real-time E-field optimization with per-trial Bayesian inference yields motor representation localization that is both efficient and precise. Beyond efficiency, the probabilistic formulation provides an interpretable measure of localization confidence, a principled stopping rule, and a natural mechanism for incorporating anatomical priors on the probable location of the cortical source. Validation against an independent ground truth such as direct electrical stimulation remains a necessary step toward clinical adoption. The limitations identified above point to concrete directions for improvement. Once validated, the method may provide a path toward more efficient presurgical motor mapping and could potentially contribute to reductions in craniotomy size, operative risk, and patient burden.

## Acknowledgements

We thank the Aalto Science-IT project for providing computational resources. We also thank Ilkka Rissanen for the manufacturing of the multi-locus TMS coil arrays and the related testing. This work was supported by the Häme Regional Fund of Finnish Cultural Foundation and by the European Research Council (ERC) under the European Union’s Horizon 2020 research and innovation programme (Grant agreement No. 810377). SP has received funding from Jenny and Antti Wihuri Foundation, Ella and Georg Ehrnrooth Foundation, KAUTE Foundation, and Finnish Foundation for Technology Promotion. IG has received funding from the Swedish Cultural Foundation, the Finnish Foundation for Technology Promotion, and the Paavo Koskinen fund from the Finnish Cultural Foundation. AMS has received funding from Jenny and Antti Wihuri Foundation and Finnish Cultural Foundation. RHM has received funding from Jenny and Antti Wihuri Foundation (Grant No. 00250241) and Tandem Industry Academia (Decision No. 366). VHS has received funding from the Research Council of Finland (Decision No. 349985). Ole Numssen and Konstantin Weise were supported by the Federal Ministry of Research, Technology and Space (Bundesministerium für Forschung, Technologie und Raumfahrt, BMFTR, Grant no. 01GQ2201 to Thomas Knösche).

## Author contributions

ML: Conceptualization, Data curation, Formal analysis, Funding acquisition, Investigation, Methodology, Project administration, Resources, Software, Validation, Visualization, Writing – original draft, Writing – review and editing. TPM: Conceptualization, Funding acquisition, Methodology, Project administration, Supervision, Writing – review and editing. SP: Funding acquisition, Investigation, Resources, Writing – review and editing. IG: Funding acquisition, Investigation, Methodology, Writing – review and editing. ON: Conceptualization, Writing – review and editing. KW: Conceptualization, Writing – review and editing. MS: Methodology, Software, Writing – review and editing. AMS: Funding acquisition, Investigation, Software, Writing – review and editing. RHM: Investigation, Writing – review and editing. VHS: Funding acquisition, Methodology, Resources, Software, Writing – review and editing. TRK: Conceptualization, Funding acquisition, Writing – review and editing. RJI: Conceptualization, Funding acquisition, Project administration, Supervision, Writing – review and editing.

## Data availability

The *ABL* Python toolkit [25] is available at https://github.com/lainem11/bayesian_localization.

