## Supplementary Material for "Adaptive Bayesian localization of motor representation areas"

### mTMS pulse delivery and strain limit

The E-field optimization returns the rate of change of current (dI/dt) $A_c$ for the $c$th coil of the array. We calibrated the mTMS device to produce the desired $A_c$, as an average dI/dt over a 60 $\mu$s rising phase of the trapezoidal pulse waveform.

Delivering pulses with the mTMS device was constrained by a maximum allowed strain on the coil set, based on the delivered current waveforms $I_c(t)$. The waveforms were simulated using a model of the power electronics [1]. The strain $S$ was calculated separately for the bottommost and topmost coils:

$$S_{\text{bottom}} = \int_t I_1(t) \left(0.9\, I_2(t) + 0.7\sqrt{I_3(t)^2 + 0.9\, I_4(t)^2} + 0.2\, I_5(t)\right),$$
$$S_{\text{top}} = \int_t I_5(t) \left(0.2\sqrt{0.9\, I_1(t)^2 + I_2(t)^2} + \sqrt{0.9\, I_3(t)^2 + I_4(t)^2}\right),$$

and the maximum of the two was selected as the pulse's strain $S$. The maximum limit for the strain was calculated as the strain $S$ of a pulse where the 4th and 5th coils are driven with 80% of their maximum stimulator output values.

### Localization maps and model fits

Localization maps were computed for the Bayesian model and the MEP amplitude regression model, with both the adaptive and randomized protocols. Figure 2 in the main text shows the results with the Bayesian model under the adaptive protocol, and results with the other method–protocol approaches are shown in Figures S1, S2, and S3.

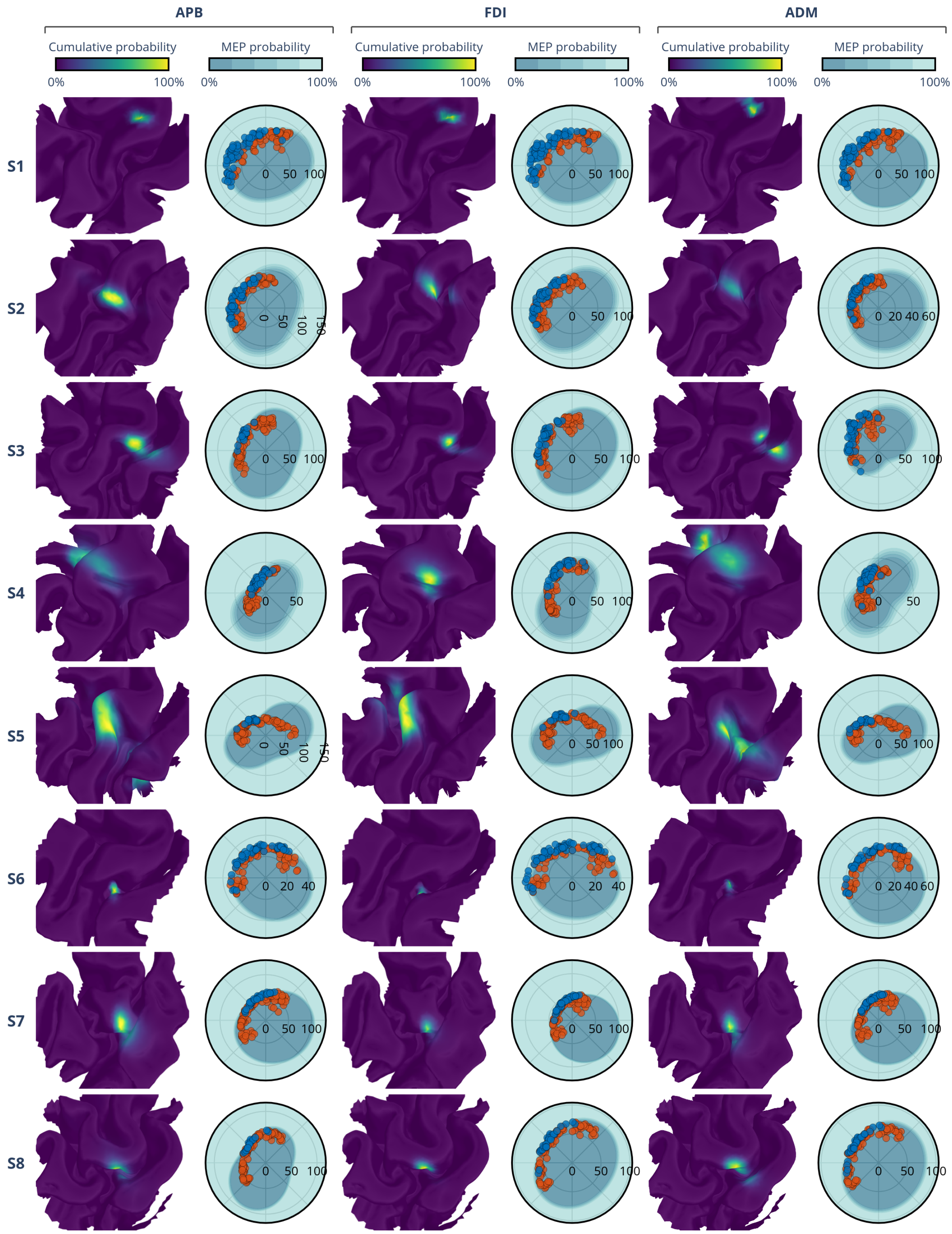


**Figure S1:** APB, FDI, and ADM localization with the Bayesian model under the randomized protocol. See Figure 2 in the main text for figure conventions.

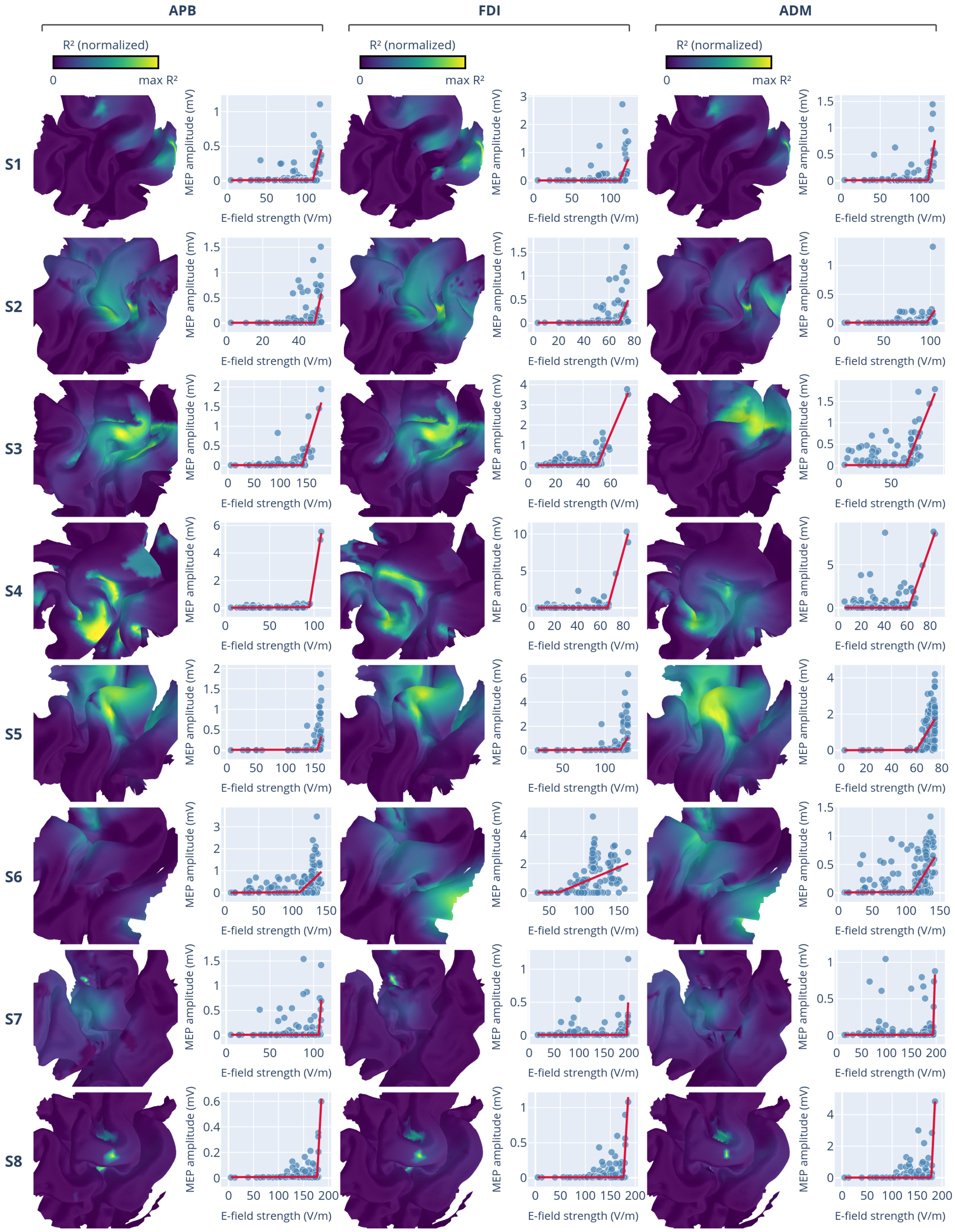


**Figure S2:** Localization with the MEP amplitude model under the adaptive protocol. Figure conventions are otherwise the same as in Figure S1, but localization maps are represented using $R^2$ and the plots on their right show the piecewise-linear regression fits at the region of highest $R^2$, with blue dots showing individual MEP amplitudes, and the red line showing the fit.

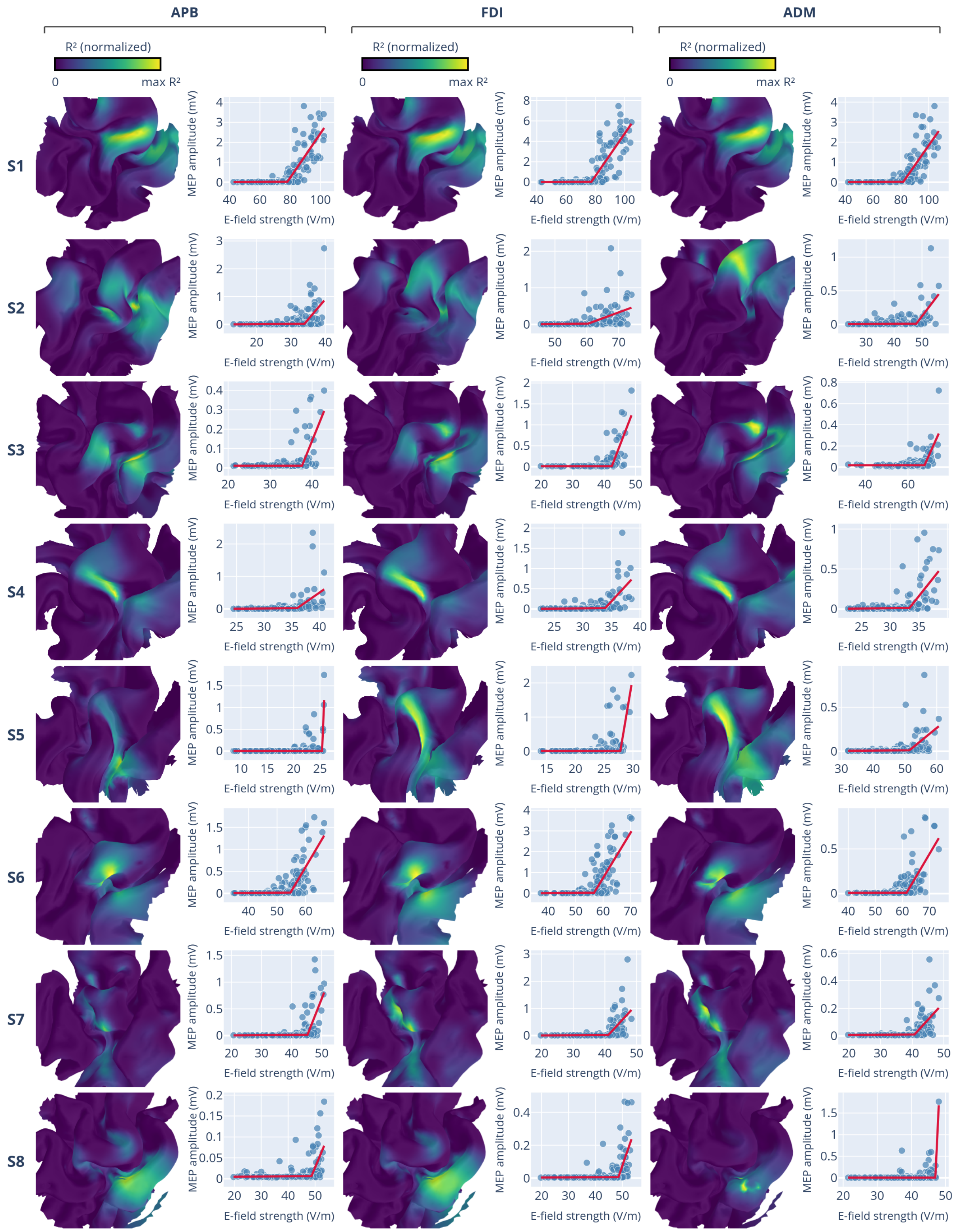


**Figure S3:** Localization with the MEP amplitude model under the randomized protocol. See Figure S2 for figure conventions.
